# Linear models replicate the energy landscape and dynamics of resting-state brain activity

**DOI:** 10.1101/2024.05.21.595246

**Authors:** Yuki Hosaka, Takemi Hieda, Kenji Hayashi, Koji Jimura, Teppei Matsui

## Abstract

The spatiotemporal dynamics of resting-state brain activity can be characterized by switching between multiple brain states, and numerous techniques have been developed to extract such dynamic features from resting-state functional magnetic resonance imaging (fMRI) data. However, many of these techniques are based on momentary temporal correlation and co-activation patterns and merely reflect stationary, linear features of the data, suggesting that the non-stationary dynamic features extracted by these techniques may be misinterpreted. To examine whether such misinterpretations occur when using techniques that are not based on momentary temporal correlation or co-activation patterns, we addressed Energy Landscape Analysis (ELA), a statistical physics-inspired method that was designed to extract multiple brain states and dynamics of resting-state fMRI data. We found that the ELA-derived features were almost identical for real data and surrogate data suggesting that the features derived from ELA were accounted for by stationary and linear properties of the real data rather than non-stationary or non-linear properties. To confirm that surrogate data were distinct from the real data, we replicated a previous finding that some topological properties of resting-state fMRI data differed between the real and surrogate data. Overall, the present finding that the energy landscape and activity dynamics of resting-state fMRI data were well captured by stationary and linear surrogate data supports the notion that linear models sufficiently describe the dynamics of resting-state brain activity.

## Introduction

Brain activity in the resting state, as measured using functional magnetic resonance imaging (fMRI) has been widely investigated for its potential applications in the non-invasive diagnosis of neuropsychiatric and neurological disorders (Fox and Raichle 2007). A common assumption regarding the dynamics of resting-brain activity is that it can be explained by transitions between multiple brain states (Vidaurre, Smith, and Woolrich 2017; Noro et al. 2022; Hutchison et al. 2013; Calhoun et al. 2014; Preti, Bolton, and Van De Ville 2016). Recent studies have reported that dynamic features (e.g., brain states) extracted from measured resting-state brain activity can better explain subject-specific phenotypes (e.g., cognitive performance) than static features (Cabral et al. 2017; Liégeois et al. 2019), suggesting the potential importance of dynamic features for applications such as diagnosis of neuropsychiatric disorders.

However, it remains an open question whether the presence of multiple brain states is supported by resting-state fMRI data. Recent statistical examinations of common analysis techniques used to extract possible brain states from resting-state fMRI data, such as sliding-window correlation analysis (Hutchison, et al. 2013; Matsui, Murakami, and Ohki 2018b) or co-activation pattern analysis (Liu, Chang, and Duyn 2013), reported that these potential brain states can be fully reproduced with surrogate data which only have a single state by construction (Laumann et al. 2016; Liégeois et al. 2017; Matsui et al. 2022). Using surrogate data, these studies extensively examined the results obtained with sliding-window correlation analysis or co-activation pattern analysis (e.g. brain states, transition probability). These surrogate data were designed to retain selected statistical properties of the real fMRI data, such as covariance structure and autocorrelation, and were produced using stationary and linear models. Crucially, these methods produced almost identical results for real data and surrogate data, which contradicted the assumptions of sliding-window correlation analysis and co-activation pattern analysis (Laumann, et al. 2016; Liégeois, et al. 2017; Matsui, et al. 2022). Thus, features extracted by sliding-window analysis or co-activation pattern analysis reflect stationary, linear properties of the real fMRI data, indicating that these analyses cannot be regarded as evidence of the non-stationarity, or multiple brain-states, of resting brain dynamics.

Given that methods based on sliding-window correlations and co-activation patterns are unable to extract dynamic features of resting-state fMRI data, a natural choice is to use alternative methods that do not use sliding-window correlations or co-activation patterns. Among these alternative methods, Energy landscape analysis (ELA) is a widely used approach inspired by statistical mechanical techniques developed for the analysis of Ising spins (Ezaki et al. 2017; Watanabe et al. 2014). On the basis of the maximum entropy principle, ELA recovers the energy landscape of resting-brain activity from the fMRI data, whose local minima correspond to the basins of attraction (i.e., brain states). Using the extracted energy landscape, ELA describes the dynamics of brain activity as transitions between brain states. Recent studies have reported that subject-level information (e.g. psychiatric conditions and cognitive scores) is reflected in the transition patterns among the states extracted by ELA (Watanabe and Rees 2017; Kang et al. 2021). Although the results of ELA could be interpreted with the view that resting-brain dynamics are non-stationary, it is unknown whether ELA captures the non-stationarity statistical properties of resting-brain activity data. In the present study, we addressed this question by applying ELA to various types of surrogate data.

Figure 1 illustrates the approach used in the present study. We first prepared real fMRI data of resting-state brain activity. ELA was applied to these data, yielding energy landscapes and transition probability matrices (a path indicated by blue arrows in Fig. 1). Next, we generated surrogate data using real fMRI data and applied the same ELA to the surrogate data (a path indicated by green arrows in Fig. 1). The surrogate data retained selected statistical properties, such as covariance structure, of the real fMRI data and were Gaussian and linear by construction. Finally, we compared the results of ELA obtained with the real data with those obtained with surrogate data (brown bidirectional arrow in Fig. 1). Any difference between the two results could be attributed to statistical properties of the real data that were not used in the surrogate data, non-Gaussianity, non-linearity of the real data, or any combination of these factors.

**Figure 1.**
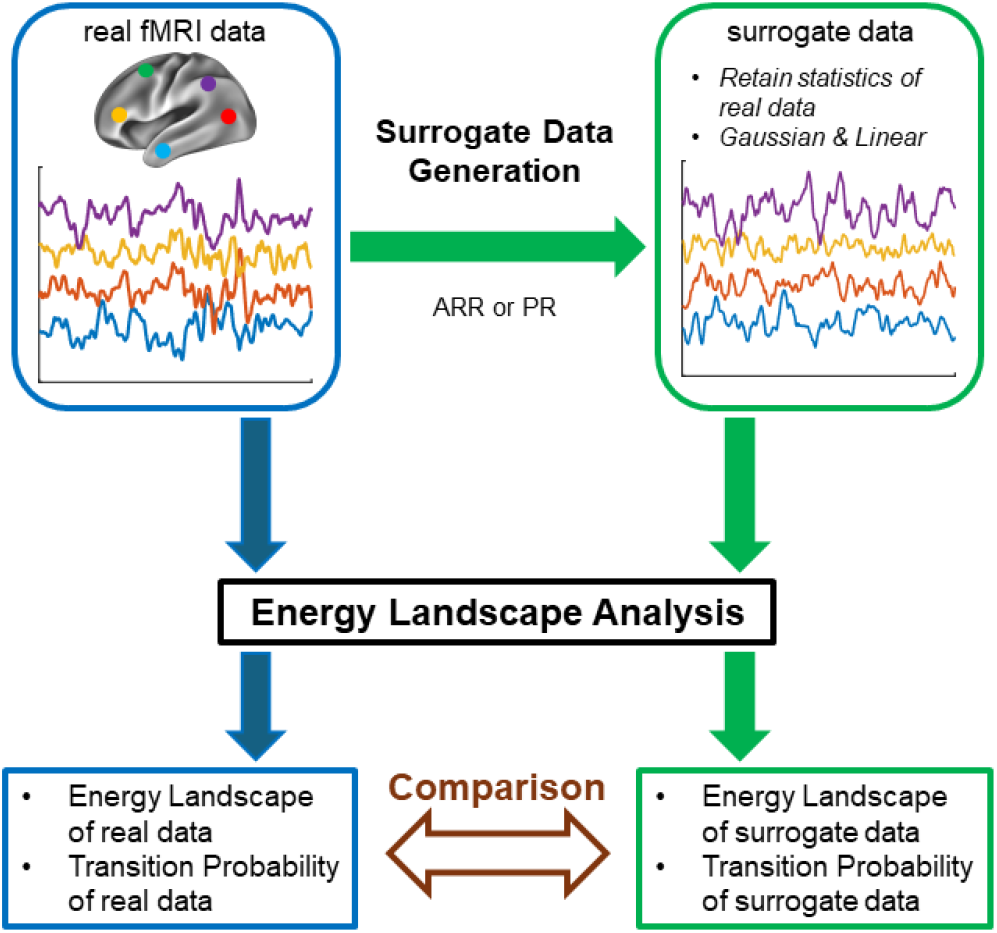
Schematics of surrogate data analysis. This schematic figure describes the strategy of the examination of ELA using surrogate data. The path indicated by blue arrows describes an ELA analysis of real resting-state fMRI data. The path indicated by green arrows describes an ELA analysis of surrogate data. In the first step of this path, surrogate time courses were constructed from the real fMRI time courses using autoregressive randomization (ARR) or phase randomization (PR). ELA was then applied to the surrogate data. Finally, in the step indicated by the brown bidirectional arrow, the results of the ELA (energy landscapes and transition probabilities) obtained in the two paths were compared.

## Materials and methods

### Dataset

For analyses of representative data, we used an example fMRI data provided by Ezaki *et al*. (Ezaki, et al. 2017). The data consisted of time courses from seven regions of interest (ROIs) within the cingulo-opercular network (CON), comprising 9,560 volumes with a repetition time (TR) of 0.72 s. For population analysis, we used the S1200 release of resting-state fMRI distributed by the Human Connectome Project (HCP; http://humanconnectomeproject.org/) (Van Essen et al. 2013). The data were preprocessed to obtain ROI-based timecourses (4000 volumes × 264 ROIs × 1002 subjects; TR, 0.72 s; see (Matsui, et al. 2022) for details). From 264 ROIs defined in (Power et al. 2011), we selected seven ROIs related to CONs, 11 ROIs related to the fronto-parietal network (FPN) and 12 ROIs related to the default mode network (DMN), whose centers were closest to the CON, FPN and DMN ROIs defined in (Fair et al. 2009). We concatenated data from two subjects, yielding 501 real data samples in total.

### Generation of surrogate data

For each sample of real data, we applied three types of linear, stationary models to generate simulated data that retained certain statistical properties of the real data (Matsui, et al. 2022; Liégeois, et al. 2017). The first model retained only the covariance structure of the real data (Static Null). Simulated data for Static Null were generated using a multivariate Gaussian distribution with covariance matrices set to those of the real data. The second model was a first-order autoregressive randomization null model (ARR). The lag of the ARR null was set to 1. Thus, ARR assumed that the fMRI data at time *t* is the sum of the linear transformation (A_*1*_) of the fMRI data at time *t-1* and zero-mean multivariate Gaussian noise with a covariance matrix (*Σ*). The parameters for the autoregressive equation (*Σ, A*_*1*_) were fitted as described previously (Liégeois, et al. 2017). Simulated data for ARR were generated using a randomly selected time point from the real fMRI data as the seed and by iteratively applying the autoregressive equation. The third model was a phase randomization null model (PR). PR retained the complete autoregressive structures of the real data as well as the covariance structures (Liégeois, et al. 2017). Simulated data for the PR null were generated by first applying a discrete Fourier transform (DFT) to the real fMRI data. Random phases were then added to the Fourier-transformed data, and inverse DFT was applied. The added phases were independently generated for each frequency but were the same across brain regions (Liégeois, et al. 2017). We referred to the simulated data produced by the null models as surrogate data.

### Energy landscape analysis (ELA)

ELA was performed as described previously (Ezaki, et al. 2017) using Matlab2023a (MathWorks, Natick, MA) with code provided by (Ezaki, et al. 2017). Briefly, for both real and surrogate data, fMRI time courses were binarized to -1 and 1. This implies that, for data with N ROIs, each volume could assume one of 2^N^ states. After binarization, a pairwise maximum entropy model was fitted to each sample of real or surrogate data, yielding an energy landscape. Basins of attraction of the energy landscapes were obtained by fitting dysconnectivity graphs (see (Ezaki, et al. 2017) for details).

### Comparison of ELA results obtained with real and surrogate data

Energy landscapes were compared by calculating Pearson’s correlation between an energy landscape of each sample of real data and an energy landscape obtained from the corresponding surrogate data.

For comparing the dynamics of real and surrogate data, we calculated a transition matrix describing the probability of switching (or staying) between basins of attraction. To obtain the transition matrix, each volume in the data was assigned to one of the basins of attraction, yielding a time course of state switching. Then, the probabilities of switching/staying from one basin to another in successive volumes were calculated. Comparisons of transition matrices were done in two methods. In the first method, we selected surrogate data with basins of attraction identical to those of the corresponding real data. An elementwise Pearson’s correlation was then calculated between the two matrices using all elements or only off-diagonal elements. In the second method, for each sample of surrogate data, we obtained the state-switching time course using basins of attraction obtained from the corresponding real data. We then calculated the transition matrix of the surrogate data and compared it with that of the real data using elementwise Pearson’s correlation (using all elements or only off-diagonal elements). Note that Pearson’s correlation using off-diagonal elements was calculated for those data that had more than three basins of attraction. For comparison of energy landscapes and transition probability, because each sample of surrogate data had a corresponding sample of real data, statistical testing was performed using a paired *t*-test.

### Topological data analysis (TDA)

We conducted Mapper-based TDA following the procedures described by Saggar and colleagues with the Matlab codes provided by the researchers (Saggar, et al. 2022). Briefly, in the first step, high-dimensional input data were embedded in a two-dimensional space using a filter function. To capture the intrinsic geometry of the data, we used a nonlinear filter function on the basis of neighborhood embedding. Specifically, Euclidian distances were calculated between all pairs of volumes. A k-nearest neighbor graph was then constructed using all volumes and calculated distances. Using the k-nearest neighbor graph, geodesic distances were calculated between all volumes in the input space. The geodesic distance was then embedded into a two-dimensional Euclidian space using multi-dimensional scaling. In the second step, overlapping two-dimensional binning was performed for data compression and noise reduction. Based on the previous study by Saggar *et al*. (Saggar, et al. 2022), we chose a resolution parameter of 14. In the third step, partial clustering within each bin was performed. Finally, a shape-graph was generated by connecting nodes from different bins when any volumes were shared by the bins.

We randomly selected 100 HCP participants to match the sample size reported in the previous study (Saggar, et al. 2022). For each participant, the time courses were concatenated across all four sessions. For statistical comparison, as in the previous study (Saggar, et al. 2022), we calculated the proportion of high-degree nodes (degree>20). The statistical significance of the difference across real and surrogate data was assessed using one-way analysis of variance (ANOVA). Note that we applied the TDA using the same fMRI data and the same procedure for generating surrogate data as we used in the ELA. Thus, any difference between ELA and TDA could not be attributed to the difference in the data, the preprocessing procedures, or the generation of surrogate data.

### Data and code availability

The data used in this study are available from the website of (Ezaki, et al. 2017) or from HCP. Code for reproducing essential results will be made available for download upon publication of the manuscript at https://github.com/teppei-matsui/EL. All codes used for the analysis will be provided upon reasonable request to the corresponding author.

## Results

### Energy landscape reflects the covariance structure of resting-brain activity

First, we compared the energy landscapes obtained with real data and surrogate data constructed by stationary null models. Figure 1a shows the energy landscapes of example real data used in a previous study (Ezaki, et al. 2017) and the surrogate data constructed from it. This example demonstrates that the energy landscapes of the real and surrogate data were highly correlated and almost perfectly overlapped for all the null models tested (Static Null, R = 0.988; ARR, R = 0.975; PR, R = 0.982) (Fig. 2a). These results indicate that linear and stationary models taking into account the second-order statistics of the data are enough to capture the shape of the energy landscape.

**Figure 2.**
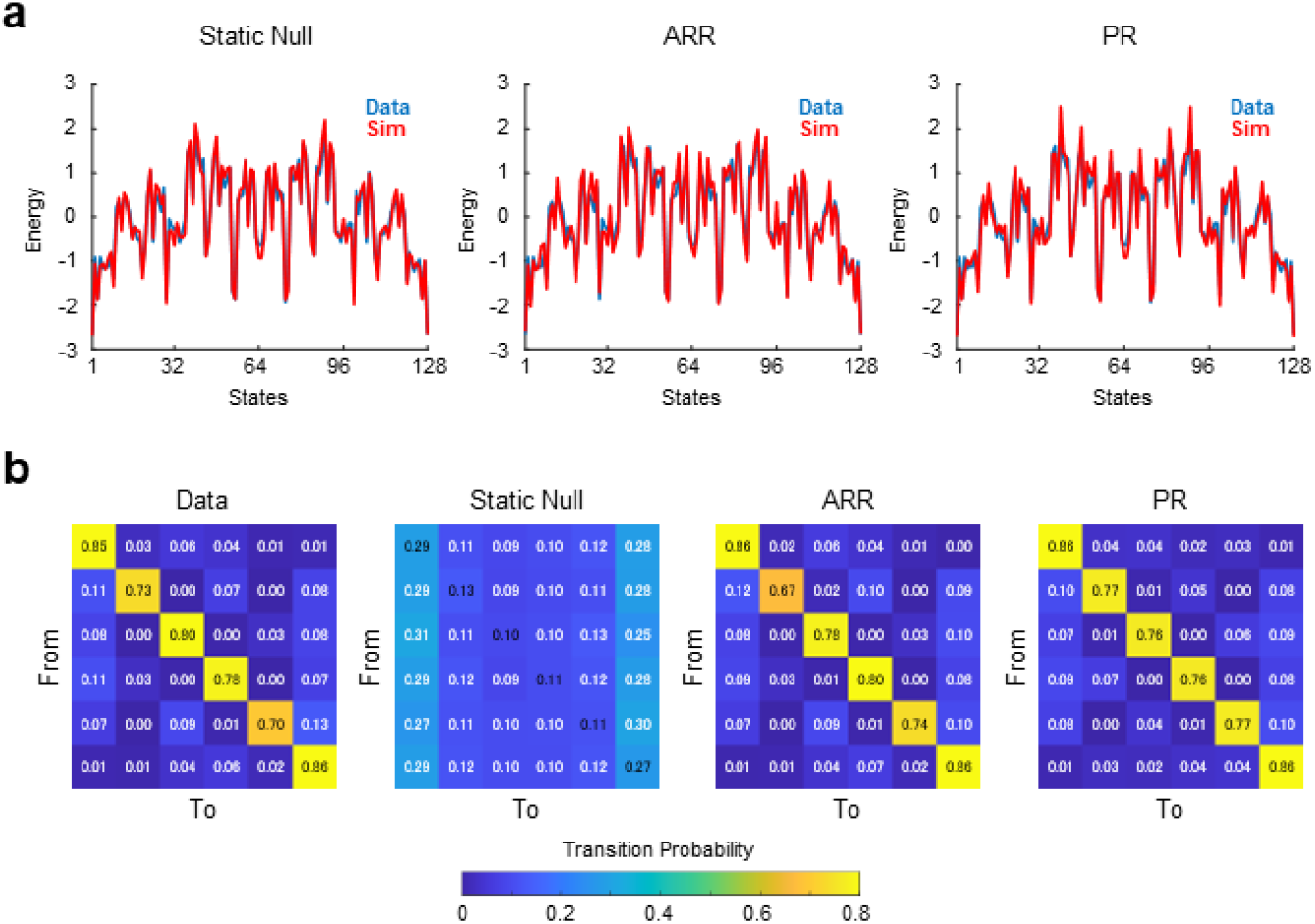
Energy landscapes and transition probability matrices for example and surrogate data calculated from it. **a**. Plots of energy landscapes. Energy landscapes for real data (blue) and surrogate data (red) are overlaid for each type of surrogate data. **b**. Matrices of transition probability between energy minima (basins). Note that energy minima were identical for the real data and all surrogate data

### Transition patterns among states reflect the autocorrelation of resting-brain activity

Next, we examined whether the transition patterns among the energy minima could be captured by the null models. We found that the basins of attraction in the energy landscapes were identical for the real and surrogate data. Figure 2b shows the transition probability matrices describing the state-switching dynamics of the real and surrogate resting brain activities. Unlike the shape of the energy landscape, transition probabilities obtained with Static Null showed low correlations with those obtained with the real data (R = 0.107). In contrast, the transition probabilities obtained with ARR and PR showed extremely high correlations with the real transition probabilities (ARR, R = 0.998; PR, R = 0.999). The correlation between transition matrices was higher for ARR and PR than for Static Null even when excluding diagonal elements (Static Null, R = 0.618; ARR, R = 0.961; PR = 0.886). The relatively high positive correlation for Static Null was likely due to the relative frequency of states reflected in the covariance structure. These results indicate that the dynamics of the transitions between energy minima can be effectively captured by linear autoregressive models.

### Test of reproducibility in a large database

To confirm whether these observations also hold in other datasets, we compared energy landscapes of real and surrogate data using a publicly available large-scale database of resting-state fMRI provided by HCP. From this database we obtained 501 samples of CON activities. Correlation coefficients of energy landscapes of the real and surrogate data were high for all tested null models (Fig. 3a). Although the differences were small, the correlation values were the highest for Static Null and lowest for ARR [Static Null, 0.871 ± 0.068 (mean ± s.d.); ARR, 0.818 ± 0.085; PR, 0.864 ± 0.070; *p* < 0.10^−10^ (uncorrected) for all pairwise comparisons, paired *t*-test]. These results suggest that the shape of the energy landscape of resting-state fMRI data can be largely captured by stationary and linear statistical properties.

**Figure 3.**
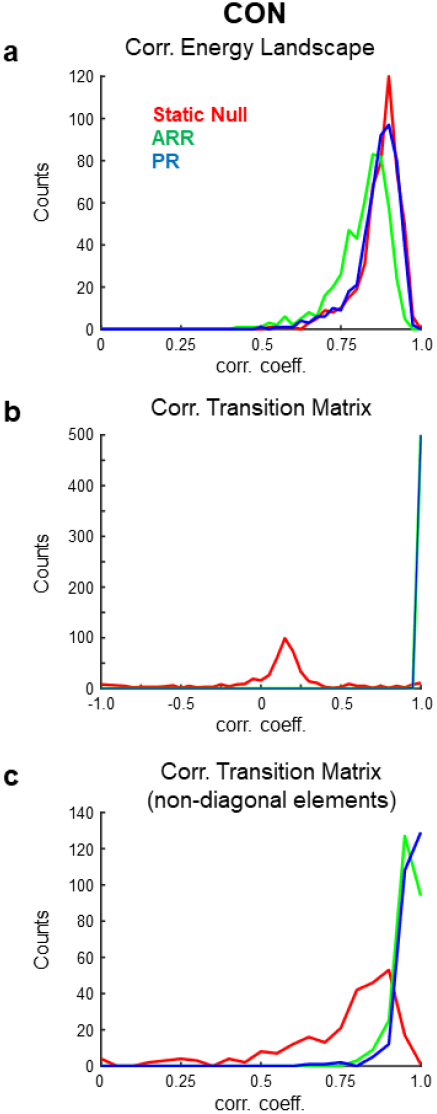
Results of population analysis based on HCP. **a**. Distribution of correlation coefficients between energy landscapes calculated from real and surrogate data. **b**. Distribution of correlation coefficients between transition matrices calculated from real and surrogate data. Note that the correlations were calculated using all elements of the transition matrices. **c**. Distribution of correlation coefficients between transition matrices calculated from real and surrogate data using only non-diagonal elements of the transition matrices.

For simulations yielding identical energy landscapes (32, 17, and 34 out of 501 simulations for Static Null, ARR, and PR, respectively), correlation coefficients between transition matrices obtained from data and simulations were high for ARR [R = 0.994 ± 0.006 (mean ± s.d.) with diagonal elements; R = 0.751 ± 0.311 without diagonal elements] and PR (R = 0.992 ± 0.011 with diagonal elements; R = 0.711 ± 0.417 without diagonal elements) but not for Static Null (R = 0.113 ± 0.125 with diagonal elements; R = 0.545 ± 0.532 without diagonal elements).

To further examine the similarity of dynamics between data and simulations, based on the high similarities of energy landscapes between them (Fig. 3a), we calculated transition probability matrices in each sample of surrogate data using the basins of attraction calculated from the corresponding real data. Transition matrices calculated with surrogate data were highly correlated with those calculated with ARR (R = 0.999 ± 0.002 with diagonal elements) and PR (R = 0.999 ± 0.003 with diagonal elements) but not with Static Null (R = 0.096 ± 0.390 with diagonal elements) (Fig. 3b). No significant difference was found between ARR and PR [*p* > 0.015 (uncorrected), paired *t*-test]. Correlations calculated only using non-diagonal elements in ARR and PR were still higher than those in Static Null (Fig. 3c), although Static Null had positive overall correlations (R = 0.747 ± 0.203 without diagonal elements). Though the difference was small, correlation values were significantly higher for PR (R = 0.956 ± 0.037 without diagonal elements) than ARR (R = 0.963 ± 0.043 without diagonal elements) [*p* < 0.004 (uncorrected), paired *t*-test]. Similar results were obtained for DMN and FPN (Fig. 4). Overall, these results confirmed the reproducibility of the observation made with the example data (Fig. 2).

**Figure 4.**
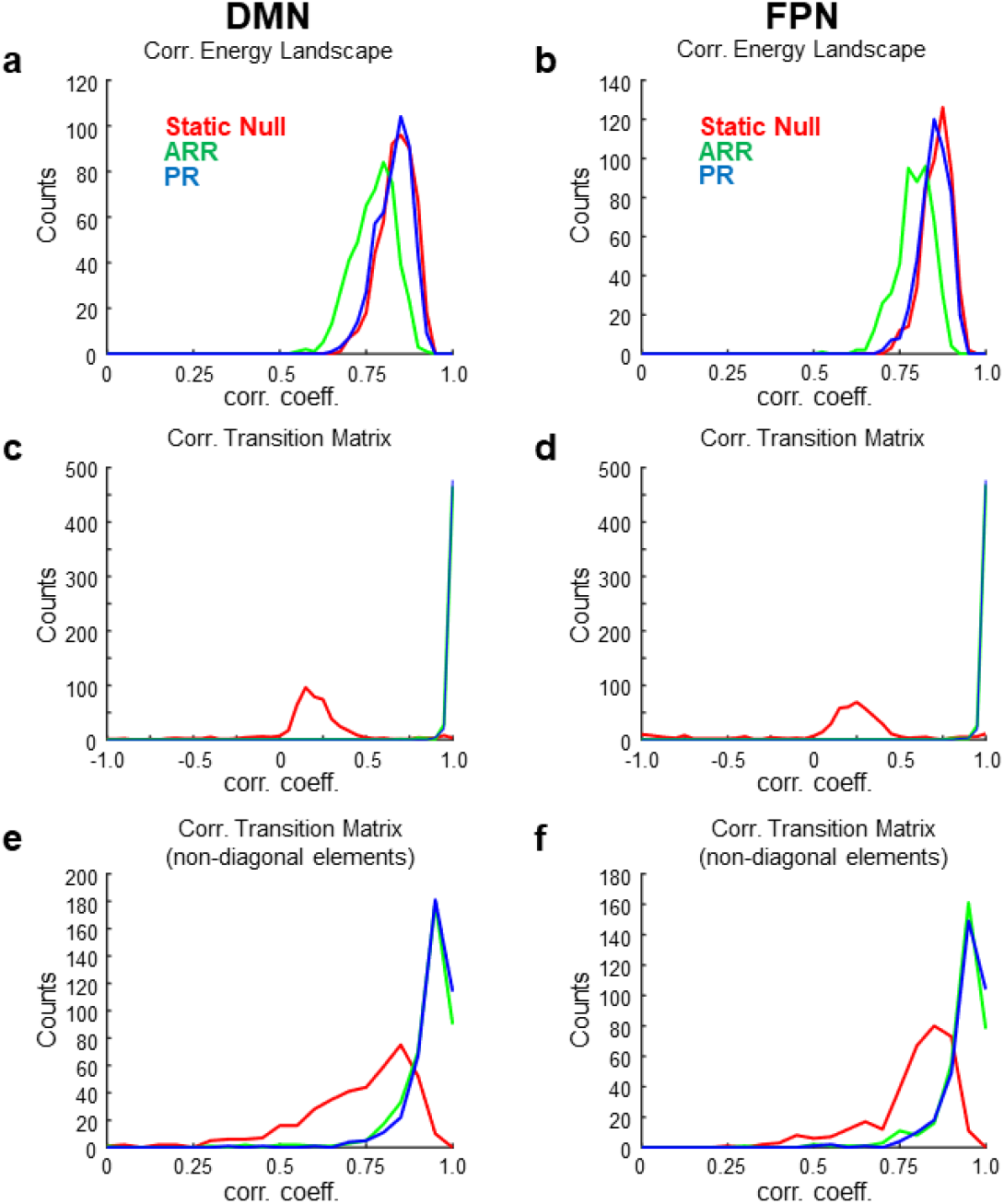
Results of population analysis based on HCP for DMN and FPN. **a**. Same as Fig.3 but for DMN and FPN. **a-b**. Distribution of correlation coefficients between energy landscapes for DMN (a) and DMN (b). **c-d**. Distribution of correlation coefficients between transition matrices for DMN (c) and FPN (d). **e-f**. Distribution of correlation coefficients between transition matrices for DMN (e) and FPN (f).

### Topological analysis confirmed the distinction between real and surrogate data

All of the results presented so far indicated that ELA yields largely identical outcomes for real resting-state fMRI data and surrogate data. This raises the possibility that real resting-state fMRI data are indeed fully describable by linear autoregressive models with residuals that have Gaussian distributions. However, previous studies reported that TDA can distinguish between real fMRI data and Gaussian, linear surrogates (Saggar, et al. 2022; Geniesse, Chowdhury, and Saggar 2022). Thus, we conducted Mapper-based TDA (Saggar, et al. 2022) to ensure that the real resting-state fMRI data contained features that were not captured by the surrogate data.

Figure 5a shows example topological landscapes of the real resting-state fMRI data and ARR and PR surrogates as visualized by Mapper-generated shape graphs (Fig. 5a). Mapper-generated shape graphs of the real data showed segregation of nodes to multiple clusters. In contrast, nodes in the shape graphs of AR and PR surrogates showed a single homogeneous cluster and appeared distinct from the real shape graphs. To quantify the difference of the graph-structure, we calculated nodal degree distributions (Fig. 5b). Compared with ARR and PR, the real data contained nodes with high degree at a higher proportion, consistent with the previous study (Saggar, et al. 2022). To test statistical significance, we assessed statistical differences in the proportion of high-degree nodes in the real versus surrogate data. Using the same threshold for high-degree node (>20) as in the previous study (Saggar, et al. 2022), we found statistically significant differences across real and surrogate data [F(2,299) = 75.49, *p* < 3.10 × 10^−27^]. These results confirmed previous reports that topological landscapes represent features of the real resting-state fMRI data that are not captured by Gaussian, linear surrogates(Saggar, et al. 2022; Geniesse, Chowdhury, and Saggar 2022). Moreover, the results suggest that the energy landscapes obtained with ELA do not capture these features.

**Figure 5.**
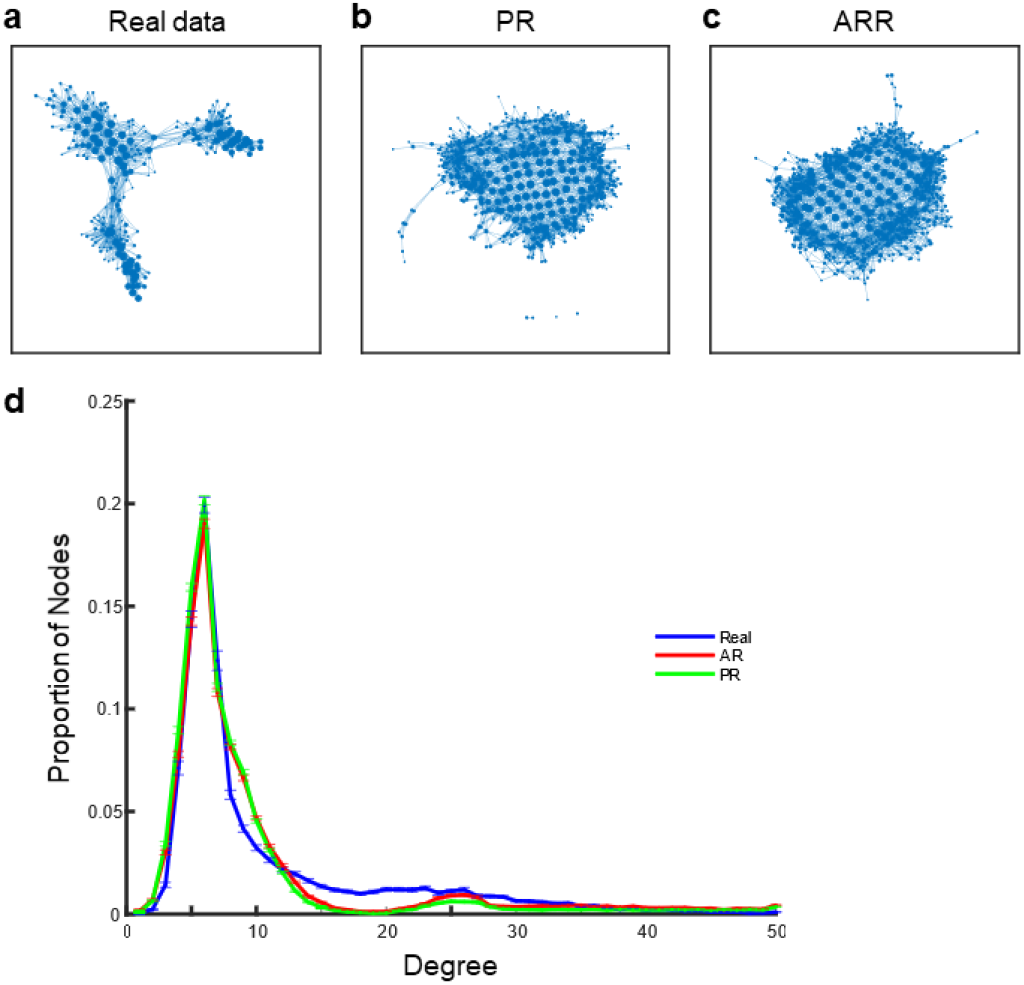
Topological landscape of real and surrogate data. **a**. Mapper shape graph of real resting-state fMRI data of an example participant. **b**. A Mapper shape graph of ARR surrogate data constructed using the real data in (a). **c**. A Mapper shape graph of PR surrogate data constructed using the real data in (a). **d**. Distributions of the degree of Mapper shape graphs across HCP participants. Error bars, SEM.

## Discussion

The current results revealed that two key results of ELA, the energy landscape and transition matrices describing the state-switching dynamics, can be explained by the stationary null models taking into account the covariance and the autocorrelation, respectively, of real resting-brain activity data. The finding that Static Null reproduced the energy landscape suggests that the energy landscape reflects the covariance structure of the resting-state fMRI data. The absence of a significant difference between ARR and PR for explaining the transition matrix suggests that first-order autocorrelation is sufficient to explain the state-switching dynamics. Thus, the present results suggest that ELA is unable to provide evidence of the non-stationarity, or multiple states, of resting brain activity.

Comparison of ELA and TDA revealed the existence of features of the real resting-state fMRI data not captured by Gaussian, linear surrogates. Consistent with previous studies (Geniesse, Chowdhury, and Saggar 2022; Saggar, et al. 2022), we found that topological landscapes could distinguish between real resting-state fMRI data and surrogate data produced by linear, gaussian models. The topological features do not necessarily reflect dynamic aspects of the data, because TDA-mapper did not use temporal information (i.e., the same topological landscapes would be obtained for temporally-shuffled data). Further characterization of the topological features obtained by TDA-mapper will be described elsewhere. It should be noted that the absence of non-linearity or non-stationarity in the features obtained with ELA does not diminish its utility in describing resting-brain activity. The present results, nevertheless, indicate that the results obtained by ELA, in particular the brain-states, should be interpreted with care [see (Matsui and Yamashita 2022) for related discussions].

From a broader perspective, the present results align with a recent proposal that macroscopic resting brain activity is best described with linear models (Nozari et al. 2024). Taken together with previous studies (Laumann, et al. 2016; Liégeois, et al. 2017; Matsui, et al. 2022), the present findings indicate that the dynamics of resting-state fMRI which resemble state-switching dynamics can be well described by simple linear models. An alternative possibility is that, because of the large amount of measurement noise in fMRI, many existing analysis methods, such as ELA, are blind to nonlinear and non-stationary features in fMRI data. To distinguish between these possibilities and determine the extent to which simple models describe macroscopic resting-brain dynamics, future animal studies using measurements with higher signal-to-noise ratio, such as calcium imaging, would be useful (Li, Ohki, and Matsui 2023; Matsui, Murakami, and Ohki 2018a).

## Conclusions

Using surrogate data analyses, we found that the features of resting-state fMRI activity extracted by ELA, namely the shape of the energy landscape and the transition patterns among the energy minima, can be largely explained by stationary and linear statistical properties of the data. This finding supports the notion that resting-state fMRI activity is well described by linear models.

## Abbreviations

ELA: Energy Landscape Analysis
CON: Cingulo-opercular Network
FPN: Fronto-parietal Network
DMN: Default Mode Network.

## Acknowledgments

Data were provided in part by the Human Connectome Project, WU-Minn Consortium (Principal Investigators: David Van Essen and Kamil Ugurbil; 1U54MH091657) funded by the 16 NIH Institutes and Centers that support the NIH Blueprint for Neuroscience Research; and by the McDonnell Center for Systems Neuroscience at Washington University.

